# Suppression of Cytotoxic T Cell Functions and Decreased Levels of Tissue Resident Memory T cell During H5N1 infection

**DOI:** 10.1101/2020.01.09.901132

**Authors:** Matheswaran Kandasamy, Kevin Furlong, Jasmine T. Perez, Santhakumar Manicassamy, Balaji Manicassamy

## Abstract

Seasonal influenza virus infections cause mild illness in healthy adults, as timely viral clearance is mediated by the functions of cytotoxic T cells. However, avian H5N1 influenza virus infections can result in prolonged and fatal illness across all age groups, which has been attributed to the overt and uncontrolled activation of host immune responses. Here we investigate how excessive innate immune responses to H5N1 impair subsequent adaptive T cell responses in the lungs. Using recombinant H1N1 and H5N1 strains sharing 6 internal genes, we demonstrate that H5N1 (2:6) infection in mice causes higher stimulation and increased migration of lung dendritic cells to the draining lymph nodes, resulting in higher numbers of virus specific T cells in the lungs. Despite robust T cell responses in the lungs, H5N1 (2:6) infected mice showed inefficient and delayed viral clearance as compared to H1N1 infected mice. In addition, we observed higher levels of inhibitory signals including increased PD1 and IL-10 expression by cytotoxic T cells in H5N1 (2:6) infected mice, suggesting that delayed viral clearance of H5N1 (2:6) was due to suppression of T cell functions *in vivo*. Importantly, H5N1 (2:6) infected mice displayed decreased numbers of tissue resident memory T cells as compared to H1N1 infected mice; however, despite decreased number of tissue resident memory T cells, H5N1 (2:6) were protected against a heterologous challenge from H3N2 virus (X31). Taken together, our study provides mechanistic insight for the prolonged viral replication and protracted illness observed in H5N1 infected patients.

**Importance:** Influenza viruses cause upper respiratory tract infections in humans. In healthy adults, seasonal influenza virus infections result in mild disease. Occasionally, influenza viruses endemic in domestic birds can cause severe and fatal disease even in healthy individuals. In avian influenza virus infected patients, the host immune system is activated in an uncontrolled manner and is unable to control infection in a timely fashion. In this study, we investigated why the immune system fails to effectively control a modified form of avian influenza virus. Our studies show that T cell functions important for clearing virally infected cells are impaired by higher negative regulatory signals during modified avian influenza virus infection. In addition, memory T cell numbers were decreased in modified avian influenza virus infected mice. Our studies provide a possible mechanism for the severe and prolonged disease associated with avian influenza virus infections in humans.

## Introduction

Influenza A viruses, members of the *Orthomyxovirus* family, cause upper respiratory infections in humans (1). Infections by seasonal influenza A virus strains (H1N1 and H3N2) are mostly self-limiting in healthy adults; however, seasonal infections can be severe in young children and the elderly (2, 3). In addition to humans, influenza viruses can infect a variety of zoonotic species including domestic poultry, pigs, horses, seals and waterfowl (4-6). Occasionally, influenza virus strains circulating in zoonotic reservoirs can cross the species barrier and cause infections in humans. Unlike seasonal H1N1 and H3N2 strains, infections with avian influenza viruses such as H5N1 and H7N9 are often severe in all age groups and cause extensive alveolar damage, vascular leakage, and increased infiltration of inflammatory cells in the lungs. The virulent nature of avian influenza viruses has been attributed to both viral and host determinants; while the viral determinants of virulence are well defined, the contribution of host responses to disease severity remain to be elucidated.

The H5N1 strain of avian influenza virus was first detected in humans during a domestic poultry outbreak in Hong Kong in 1997 (7, 8). Despite considerable efforts for containment, H5N1 strains have spread globally and are now endemic in domestic poultry on several continents. Over the past 20 years, H5N1 viruses from infected domestic poultry have crossed the species barrier, causing severe and often fatal infections in humans with mortality rates as high as 60% (9). Many of the viral components critical for the enhanced virulence of H5N1 have been identified through the generation of recombinant and/or reassortant viruses (10) (11, 12). Prior studies have shown that the multibasic cleavage site (MBS) in the viral hemagglutinin of H5N1 facilitates higher viral replication and mediates extrapulmonary spread (13-15). In addition, our group has recently demonstrated that the endothelial cell tropism of H5N1 contributes to barrier disruption, microvascular leakage, and subsequent mortality (12). Moreover, polymorphisms that increase viral replication have been identified in the viral polymerase subunits of H5N1 strains (16-20). Together, these studies have helped to define the viral components that are responsible for the enhanced virulence of H5N1.

Apart from viral determinants, overt and uncontrolled activation of the innate immune responses also contribute to the disease severity associated with H5N1 infection (21, 22). Histological analyses of lungs from fatal H5N1 cases demonstrate severe immunopathology, as evidenced by excessive infiltration of immune cells into the lungs and higher numbers of viral antigen positive cells in the lungs (23, 24). In corroboration with these studies, H5N1 viruses have been shown to induce higher DC activation and increase cytokine production as compared to H1N1 viruses (25). Moreover, studies with H5N1 strains in animal models demonstrate hyperactivation of resident immune cells in the lungs and a consequent upsurge in cytokine levels (26, 27). As such, these heightened proinflammatory responses result in the excessive recruitment of neutrophils and inflammatory monocytes into the lungs, correlating with severe disease (24). Despite robust activation of innate immune responses against H5N1 infection, higher and prolonged virus replication can be detected in the lungs of infected individuals, suggesting a possible dysregulation of adaptive immune responses(28).

We have previously demonstrated that appropriate activation of respiratory DC is required for effective T cell responses against a mouse adapted H1N1 strain (29). Here, we sought to determine if excessive activation of innate immune cells during avian H5N1 infection impairs subsequent adaptive T cell responses. In order to investigate the immune responses against H5N1 in comparison to a mouse adapted H1N1 strain, we generated a closely matched recombinant H5N1 virus carrying the 6 internal genes of H1N1 (H5N1 (2:6)). Our studies demonstrated that H5N1 (2:6) infection in mice induced higher lung DC activation and promoted increased migration of lung DC to the draining lymph nodes, resulting in increased numbers of virus specific CD8+ and CD4+ T cells in the lungs as compared to H1N1 infected mice. Despite higher numbers of virus specific T cells, we observed delayed clearance of H5N1 from the lungs, which correlated with higher PD-1 expression and increased production of the anti-inflammatory cytokine IL-10 by T cells in H5N1 infected mice. Importantly, we observed lowered numbers of virus specific tissue resident memory T cells in H5N1 infected mice as compared to H1N1 infected mice. Taken together, our study demonstrates that hyperactivation of innate immune cells during H5N1 infection impairs cytotoxic T cell functions as well as subsequent generation of influenza virus specific tissue resident memory T cells.

## Results

### H5N1 infection induces higher activation of innate immune cells

To establish if infection with a low pathogenic H5N1 virus results in higher activation of innate immune cells, we infected C57BL/6 mice with a recombinant H5N1-GFP (A/Vietnam/1203/2004) or H1N1-GFP (A/Puerto Rico/8/1934, PR8 strain) virus and measured the activation status of different cell populations in the lungs by quantifying cell surface upregulation of CD86. For comparison, we utilized the mouse adapted H1N1 (A/Puerto Rico/8/1934, PR8) strain, as it replicates efficiently in murine lungs. We observed higher upregulation of CD86 on both types of lung resident DC (CD103+ DC and CD11b+ DC) in mice infected with H5N1-GFP as compared to H1N1-GFP (Figure 1A-B). In addition, we observed higher upregulation of CD86 on inflammatory DC and inflammatory monocytes from H5N1-GFP infected mice as compared to H1N1-GFP infected mice, demonstrating that H5N1 infection results in higher activation of innate immune cells (Figure 1C-D).

**Figure 1:**
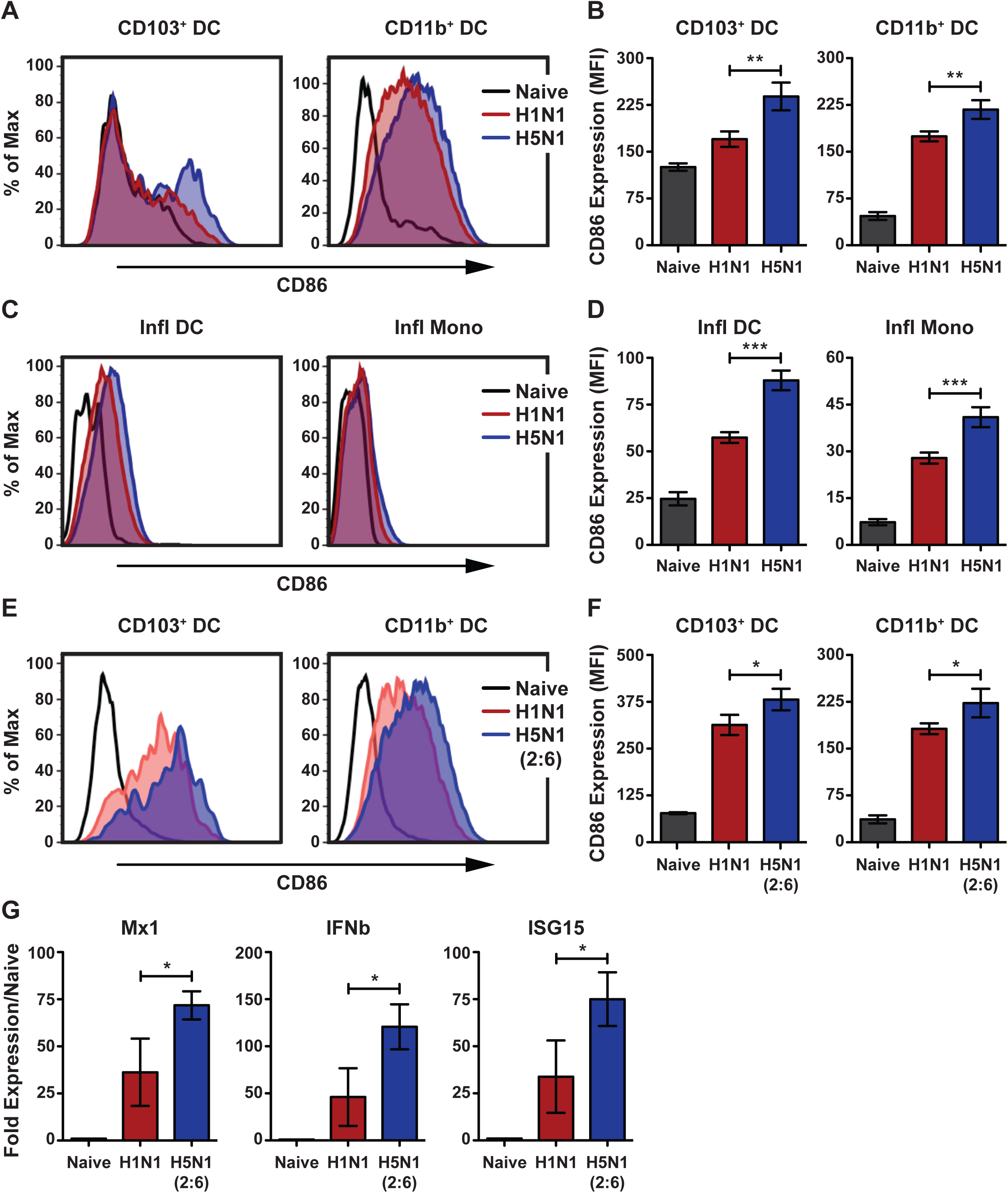
H5N1 virus stimulates higher activation of dendritic cells in the lungs. (A-D) C57BL/6 mice (n=3-4 per group) were infected with 5×10^4^ PFU of H1N1-GFP or H5N1-GFP and cell surface expression of co-stimulatory molecule CD86 was measured flow cytometry. (A) Representative histograms comparing CD86 expression on lung DC subsets. (B) Quantification of CD86 expression on lung DC subsets. CD86 expression levels are shown as mean fluorescent intensity (MFI). (C-D) Comparison of CD86 expression on inflammatory DC and monocytes. (C) Histogram plot of CD86 expression. (D) Quantification of CD86 expression. (E-F) Comparison of CD86 expression on DC subsets in mice infected with H5N1 (2:6) and H1N1. C57BL/6 mice were infected with 100 PFU of H1N1 or H5N1 (2:6) and CD86 expression was measured flow cytometry. (E) Representative histograms of CD86 expression on lung DC subsets. (F) Quantification for panel E shown as MFI. (G) Comparison of Mx1, ISG15, and IFNβ expressions between H5N1 (2:6) and H1N1 infected lungs. Total RNA was extracted from lung homogenates of infected mice isolated on day 4 pi and subjected to qRT-PCR analysis. The values are expressed as mean ± SD. *, **, *** denotes significance of <0.05, <0.01, <0.001, respectively. Data are representative of at least three independent experiments.

Next, to determine if the HA and NA of H5N1 virus are sufficient to induce higher activation of innate immune cells, we generated a 2:6 reassortant virus carrying the HA and NA from H5N1 with the 6 internal genes of PR8 (H5N1 (2:6)) and compared it to the parental strain in subsequent studies. In this way, we can minimize the differences in viral replication between H5N1 and H1N1, as well as monitor T cell responses against the same epitopes in the internal viral genes. To confirm higher activation of innate immune cells by the H5N1 (2:6) reassortant strain, C57BL/6 mice were infected with H5N1 (2:6) or H1N1 and the levels of CD86 were analyzed by flow cytometry on day 2 post-infection (pi). In mice infected with H5N1 (2:6), we observed higher expression of CD86 on both types of lung resident DC (CD103+ DC and CD11b+ DC) as compared to H1N1 infected mice (Figure 1E-F). In addition, we observed increased expression of IFNβ and interferon stimulated genes (ISG) in the lungs of H5N1 (2:6) infected mice on day 4 pi as compared to H1N1 infected mice (Figure 1G). Together, these results demonstrate that the HA and NA of H5N1 can induce higher innate immune responses in the lungs.

### H5N1 (2:6) infection stimulates increased migration of lung DC to the MLN

Upon acquisition of viral antigens and subsequent activation, lung DC upregulate CCR7 and migrate to the mediastinal lymph nodes (MLN) for priming of naïve T cells. To determine if hyperactivation of lung DC alters their migration to the lymph nodes, we infected mice with H5N1 (2:6) or H1N1 and analyzed the levels of CCR7 upregulation by flow cytometry and monitored the levels of lung DC accumulation in the MLN via CFSE labeling. We observed increased upregulation of CCR7 on the CD103+ DC subset in H5N1 (2:6) infected mice as compared to H1N1 infected mice (Figure 2A-B); however, CCR7 expression was comparable in the CD11b+ DC subset in both groups. Next, to determine the levels of lung DC migration to the MLN, we labeled cells in the respiratory tract by instilling CFSE dye on day 2 pi and measured the levels of CFSE positive lung DC in the MLN after 16h. As compared to H1N1 infected mice, we observed increased accumulation of CFSE+ lung DC in the MLN of H5N1 (2:6) infected mice (Figure 2C-E). In addition, we observed increased numbers of total lung DC in the MLN of H5N1 (2:6) infected mice as compared to H1N1 infected mice (Figure 2E). These data demonstrate that H5N1 (2:6) infection induces higher activation of lung DC, resulting in increased migration and accumulation of lung DC in the MLN.

**Figure 2:**
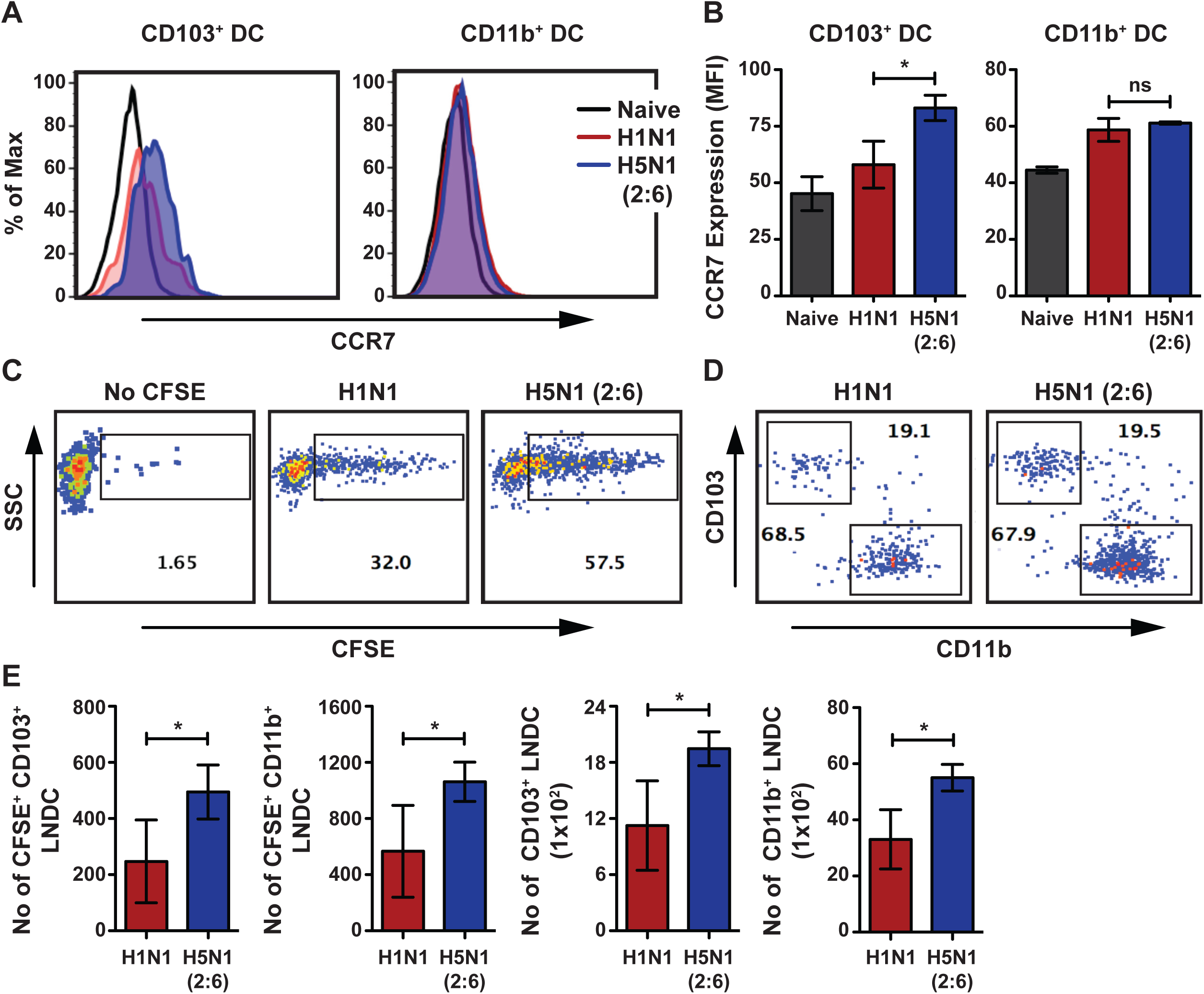
H5N1 (2:6) infection induces higher upregulation of CCR7 and migration of lung DC. C57BL/6 mice (n=3-4 per group) were intranasally infected with 100 PFU of H1N1 or H5N1 (2:6) and lung DC activation and migration was analyzed by flow cytometry. (A) Representative histogram comparing the expression of CCR7 on CD103+ DC or CD11b+ DC subsets on day 2 pi. (B) Quantification for panel A. CCR7 expression levels are shown as MFI. (C-D) C57BL/6 mice were infected with 100 PFU of H5N1 (2:6) or H1N1 and instilled with 50μl of 8mM CFSE at day 2pi. After 16h, the number of CFSE+ migratory DC present in the MLN was analyzed by flow cytometry. (C) Representative FACS plots showing CFSE+ population in the MLN. (D) Relative levels of CFSE positive CD103+ and CD11b+ DC in the MLN. (E) Bar charts showing the number of CFSE+ DC subsets in the MLN. (F) Bar chart showing total numbers of DC subsets in the MLN. The values are expressed as mean ± SD. * denotes statistical significance of <0.05; ns denotes not significant. Data are representative of at least two independent experiments.

### Mice infected with H5N1 (2:6) show robust activation of T cell responses but display delayed viral clearance

Next, we determined if the higher numbers of DC observed in the MLN of H5N1 (2:6) infected mice resulted in enhanced T cell responses and viral clearance. To evaluate primary T cell responses, C57BL/6 mice were infected with 100 PFU of H5N1 (2:6) or H1N1 and T cell responses were measured on day 8 pi by tetramer staining and by monitoring for cytokine production upon *ex vivo* stimulation. Using tetramers specific for viral NP or PA, we observed increased frequencies of both NP and PA tetramer positive CD8+ T cells in H5N1 (2:6) infected mice as compared to H1N1 infected mice (Figure 3A-B). The absolute numbers of virus specific CD8 T cells were also higher in H5N1(2:6) infected mice as compared to H1N1 infected mice (Figure 3C). In addition, *ex vivo* stimulation with X-31 (H3N2) virus or viral peptides showed increased frequencies of interferon gamma (IFNγ) and granzyme B (GrB) producing CD8+ T cells in H5N1 (2:6) infected mice as compared to H1N1 infected mice (Figure 3D). Moreover, H5N1 (2:6) infected mice showed increased frequencies of IFNγ and GrB producing CD4+ T cells as compared to H1N1 infected mice (Figure 3E). These results demonstrate that hyperactivated lung DC promote robust activation of virus specific T cell responses in the lung.

**Figure 3:**
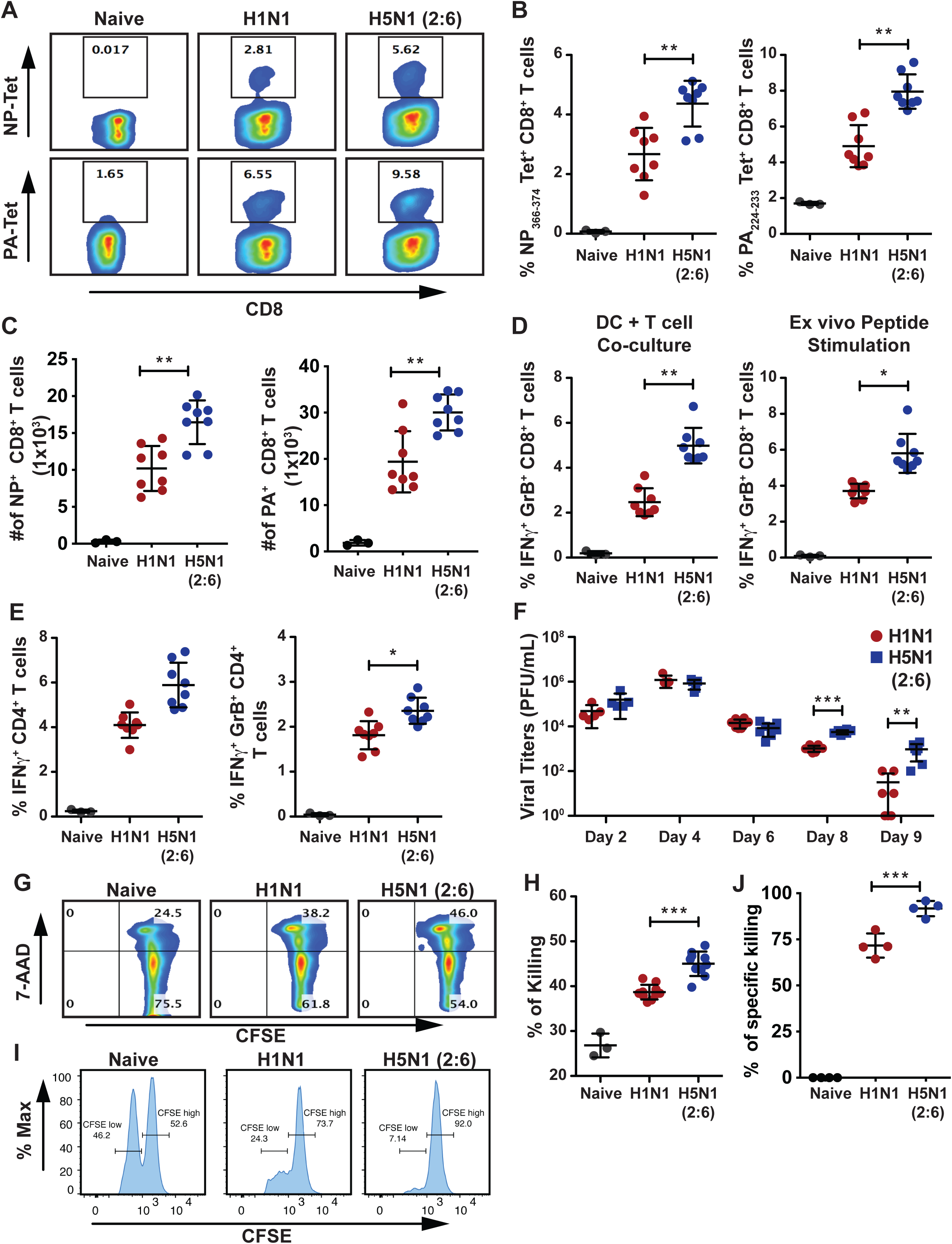
Mice infected with H5N1 (2:6) mount robust T cell responses but show delayed viral clearance. (A-E) C57BL/6 mice (n=3-4/group) were infected with 100 PFU of H1N1 or H5N1 (2:6) and on day 8 pi, T cells from the lungs were isolated and evaluated in various assays. (A-C) Comparative analysis of lung CD8+ T cells from H5N1 (2:6) or H1N1 infected mice by NP or PA tetramer staining. (A) Representative FACS plots for NP or PA tetramer staining. (B) Relative frequency of tetramer positive CD8+ T cells. (C) Absolute numbers of virus specific CD8+ T cells. (D-E) Comparative analysis of cytokine production in T cells isolated from the lungs of H5N1 (2:6) or H1N1 infected mice. T cells were co-cultured with BMDC either infected with X31 (H3N2) or pulsed with NP peptide, and the frequencies of IFNγ and GrB producing T cells were analyzed by flow cytometry. (D) Relative frequency of cytokine producing CD8+ T cells. (E) Relative frequency of cytokine producing CD4+ T cells stimulated with NP peptide. (F) Evaluation of viral loads in the lungs of infected mice. C57BL/6 mice were infected with 100 PFU of H1N1 or H5N1 (2:6) and at various times pi, viral loads in the lungs were measured by standard plaque assay. (G-H) *Ex vivo* analysis of cytotoxic T cell functions. CFSE labeled splenocytes pulsed with NP peptide were co-cultured with lung CD8+ T cells for 8hrs, followed by staining with 7-AAD. *Ex vivo* cytotoxic effects of CD8+ T cells were evaluated by analyzing 7-AAD positive splenocytes. (G) Representative FACS plots for 7-AAD positive cells and (H) relative level of killing by T cells shown as percentage of 7-AAD positive cells. (I-J) *In vivo* analysis of cytotoxic T cell functions. (I) Representative FACS plots for *in vivo* killing of adoptively transferred NP pulsed splenocytes in H1N1 or H5N1 (2:6) virus infected mice and (J) relative level of kiiling by T cells. The values are expressed as mean ± SD. *, **, *** denotes significance of <0.05, <0.01, <0.001, respectively. Data in panels A-F are from two independent experiments pooled together. Data in panel G-J are from one experiment.

In the mouse model of influenza virus, innate immune cells restrict viral replication prior to the establishment of adaptive T cell responses. However, after day 6 pi, T cells primed in the MLN migrate to the lungs and participate in the clearance of virus infected cells. Therefore, we determined if the higher numbers of virus specific T cells observed in H5N1 (2:6) infected mice resulted in efficient viral clearance in the lungs. C57BL/6 mice were infected with 100 PFU of H5N1 (2:6) or H1N1, viral loads in the lungs were measured by plaque assay at various days pi. Prior to and including day 6 pi, we observed similar viral loads in the lungs of both groups of infected mice, suggesting that both viruses replicate to similar levels (Figure 3F). However, on day 8 and day 9 pi, we observed higher viral loads (∼5-10 fold) in the lungs of H5N1 (2:6) infected mice as compared to H1N1 infected mice. These results demonstrate that, despite the presence of more virus specific T cells in the lungs, viral clearance was delayed in H5N1 (2:6) infected mice.

To understand the basis for the delayed clearance of H5N1 (2:6) in the lungs, we evaluated the functionality of T cells by *in vitro* T cell killing assay. In this assay, T cells isolated from H5N1 (2:6) or H1N1 infected mice were co-cultured with CFSE labeled splenocytes pulsed with NP peptide, and the amount of target cell death was determined by quantification of 7-AAD positive splenocytes. Interestingly, we observed more splenocyte death in co-cultures containing T cells from H5N1 (2:6) infected mice as compared to co-cultures containing T cells from H1N1 infected mice (Figure 3G-H). Next, we performed *in vivo* killing assay with peptide pulsed splenocytes. Splenocytes were labeled with either low CFSE or high CFSE and pulsed with influenza virus NP peptide or control peptide, respectively. Splenocytes were adoptively transferred into mice previously infected with either H5N1(2:6) or H1N1 (day 8pi; Figure 3I-J) and 8hrs post adoptive transfer mice splenocytes were analysed for CFSE+ cells. Our data demonstrate that cytotoxic T cells from H5N1 (2:6) infected mice can effectively kill peptide pulsed splenocytes both *in vitro* and *in vivo*.

### Cytotoxic T cells from H5N1 (2:6) infected mice show higher expression of PD1 and IL-10

T cell functions can be modulated by stimulatory as well as inhibitory signals. Prior studies demonstrate that during influenza virus infection, T cell functions can be suppressed by PD-1/PD-L1 interactions and by the anti-inflammatory cytokine IL-10(30-32). In addition, PD-1 has shown to be upregulated in T cells in response to direct activation of TCR. Although the T cells isolated from H5N1 (2:6) infected mice were efficient in killing peptide pulsed splenocytes, we observed delayed viral clearance in the lungs (Figure 3F-G). Thus, we investigated if the T cell functionality was suppressed *in vivo* through PD-1/PD-L1 interactions by measuring the expression of PD-1/PD-L1 by flow cytometry. We observed significantly higher levels of PD-1 on CD8+ T cells isolated from H5N1 (2:6) infected mice as compared to H1N1 infected mice (Figure 4A-B). Next, we analyzed different cellular compartments in the lungs for PD-L1 expression and observed significantly higher levels of PD-L1 on inflammatory monocytes (CD11b+ Ly6C^hi^ Ly6G-) isolated from H5N1 (2:6) infected mice as compared to H1N1 infected mice (Figure 4C). In addition, we observed increased numbers of inflammatory monocytes in H5N1(2:6) infected mice group (Figure 4D). However, the levels of PD-L1 on other cellular compartments in the lungs including inflammatory DC were similar between the two groups. Next, we measured IL-10 production in T cells isolated from infected mice to determine the possibility of IL-10 mediated suppression of T cell functions. T cells isolated from H5N1 (2:6) or H1N1 infected mice on day 8 pi were co-cultured with DC pulsed with MHC-I or MHC-II peptide or infected with X-31 (H3N2) virus, and production of IFNγ and IL-10 in T cells was measured by flow cytometry. We observed increased production of IFNγ and IL-10 in both CD8+ and CD4+ T cells isolated from H5N1 (2:6) infected mice as compared to H1N1 infected mice (Figure 4E-F). Taken together, these results demonstrate that H5N1 (2:6) infection results in higher expression of inhibitory signals such as PD-1 and IL-10 by T cells, which likely suppress cytotoxic T cell functions *in vivo*.

**Figure 4:**
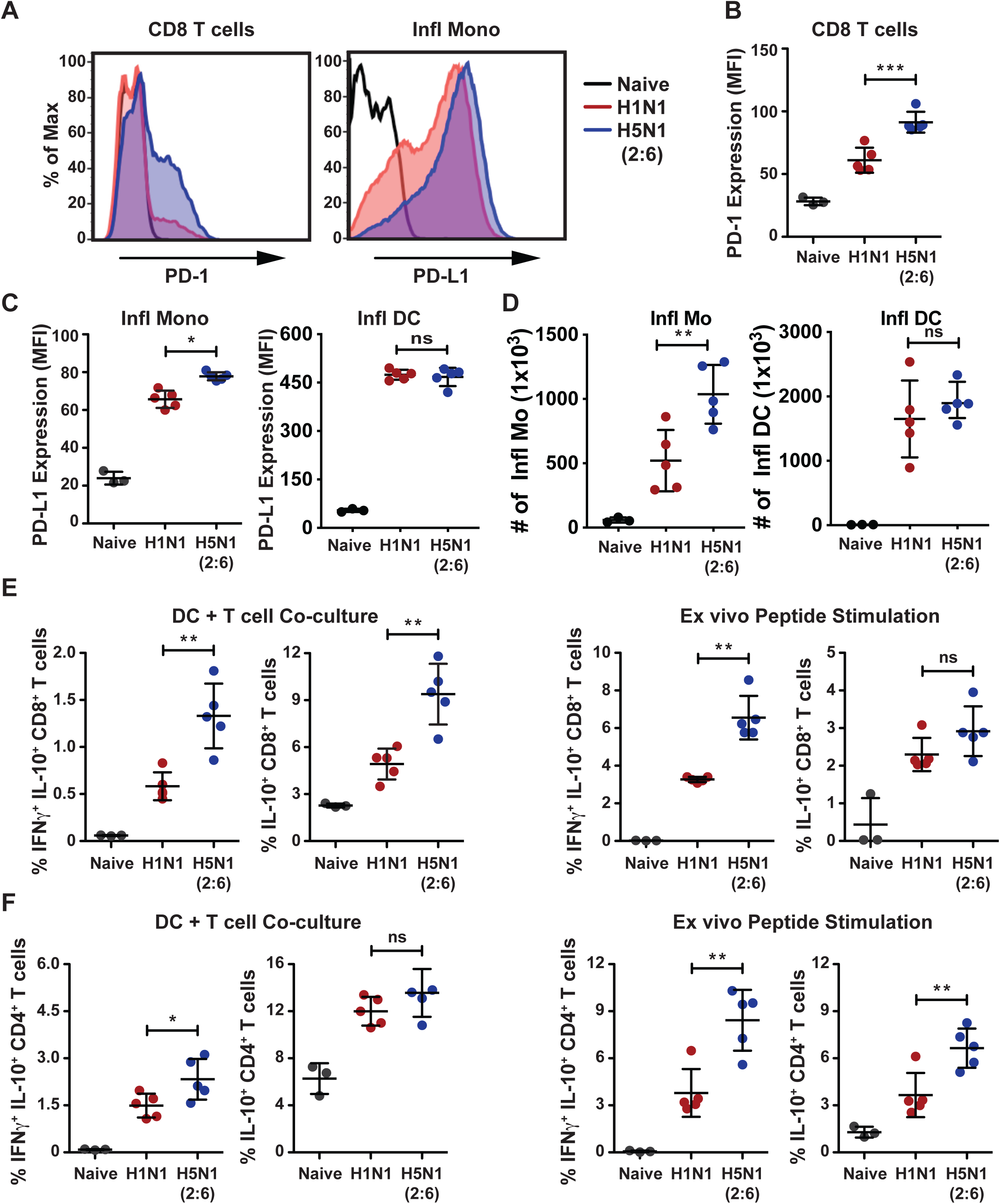
H5N1 (2:6) infection induces higher expression of PD-1 and IL-10 in cytotoxic T cells. C57BL/6 mice were infected with H5N1 (2:6) or H1N1 virus and on day 8pi, expression of PD-1 and production of IL-10 in CD8+ T cells were measured *ex vivo* upon co-culture with infected DC or peptide pulsed DC by flow cytometry. PD-L1 expression on inflammatory monocytes was also measured by flow cytometry. (A) Representative histograms showing expression of PD-1 on CD8+ T cells and PD-L1 on Ly6C^+^ inflammatory monocytes. (B) quantification for PD-1 expression in CD8 T cells as MFI. (C) Quantification of PD-L1 expression in inflammatory monocytes and inflammatory DCs. (D) Absolute numbers of inflammatory monocytes and inflammatory DCs. (E) Quantification of IL-10 producing CD8+ T cell frequencies in X-31 infected DC-T cell co-culture (upper panel) and NP peptide pulsed DC-T cell co-culture (lower panel). (E) Cytokine production in CD4+ T cells. Frequencies of IFNγ and IL-10 or IL-10 alone producing CD4+ T cells in X-31 infected DC-T cell co-culture (left panels) and NP peptide pulsed DC-T cell co-culture (right panels). The values are expressed as mean ± SD. *, **, *** denotes significance of <0.05, <0.01, <0.001, respectively.

### H5N1 infected mice show decreased numbers of memory T cells in the lung parenchyma

Upon clearance of viral infection, a portion of virus specific T cells differentiate into tissue resident memory T (T_RM_) cells, which play an important role in providing heterosubtypic immunity against subsequent influenza virus infections(33). As we observed higher upregulation of inhibitory signals (PD-1 and IL-10) on T cells from H5N1 (2:6) infected mice, we investigated if T_RM_ responses were also impaired. On day 30 pi, we analyzed the lung parenchyma for memory T cells that exhibit the T_RM_ phenotype (CD69+ CD44+ CD103+) by flow cytometry (34, 35). Circulating T cells were excluded by intravenous injection of labelled anti CD8β antibody prior to euthanizing mice and excluding this population from analysis (Figure S2). We observed lowered numbers of NP tetramer positive CD8+ T_RM_ cells in the lungs of H5N1 (2:6) infected mice as compared to H1N1 infected mice (Figure 5A). Similarly, the numbers of CD4+ T_RM_ cells were lowered in H5N1 (2:6) infected mice as compared to H1N1 infected mice (Figure 5B). These results demonstrate that H5N1 (2:6) infection results in decreased differentiation of lung resident memory T cells. Next, to determine if the decreased numbers of tissue resident memory cells affect protection from future challenge, C57BL/6 mice were infected with 50PFU of H1N1 or H5N1(2:6) virus and subsequently challenged with a heterologous H3N2 strain (X-31), a reassortant strain that share 6 internal genes with H1N1 and H5N1(2:6) viruses. We did not observe significant differences in weight loss between H1N1 and H5N1(2:6) infected groups upon lethal challenge with the H3N2 (X-31) strain. These data suggest that the lowered levels of memory T cells in H5N1(2:6) infection does not impact protection against heterologous strains.

**Figure 5.**
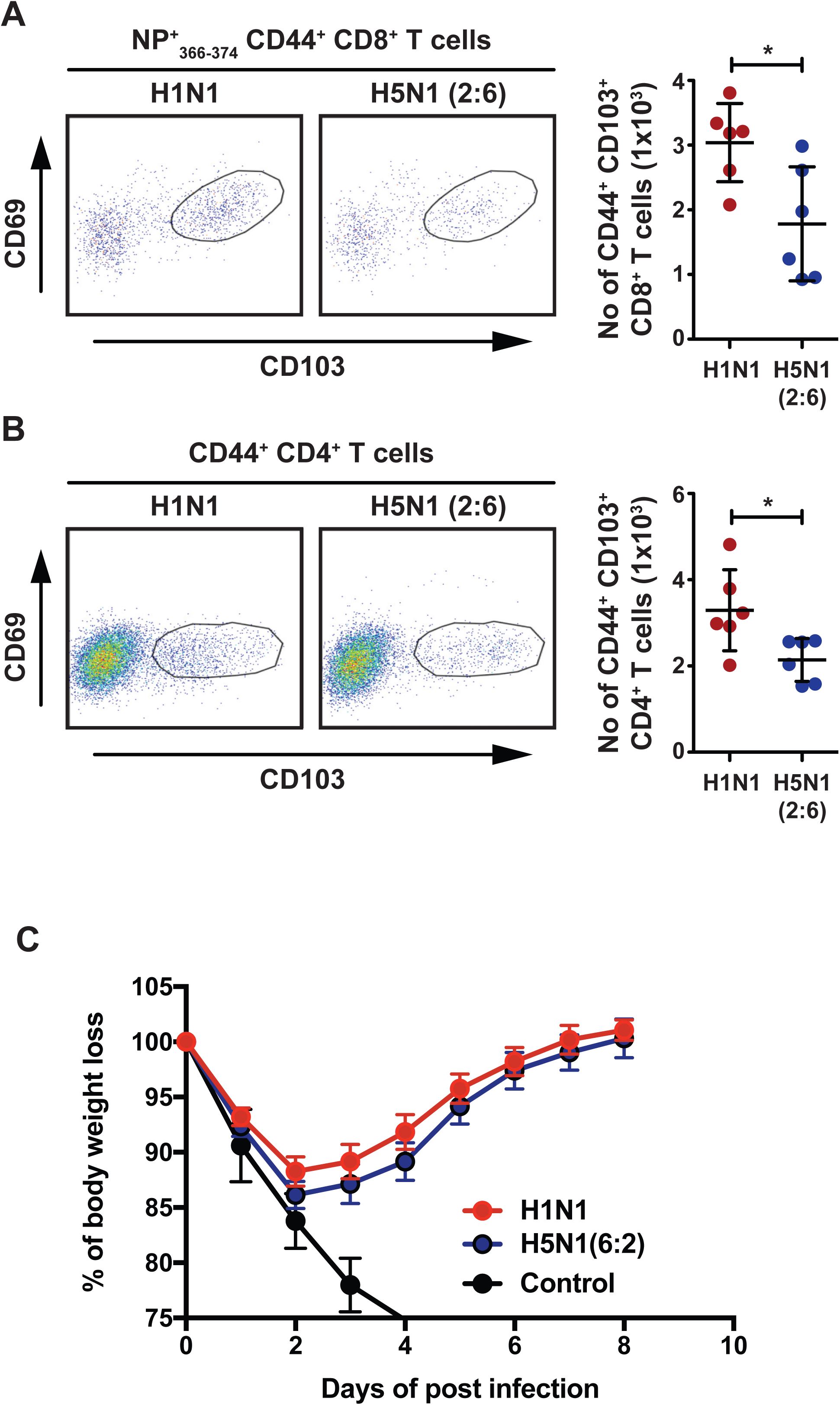
H5N1 (2:6) infection results in decreased numbers of tissue resident memory T cells in the lung parenchyma. C57BL/6 mice (n=3) were infected with H5N1 (2:6) or H1N1 virus and on day 30 pi, the frequency and absolute number of lung resident memory cells was analyzed by flow cytometry. (A) Lung resident memory CD8+ T cell responses. Representative FACS plots for lung resident memory CD8+ T cells (gated on NP+ _366-374_ CD44+ CD8α+ CD8β-T cells) that display CD44+ CD69+ CD103^hi^ phenotype (left) and the absolute numbers of tissue resident memory CD8+ T cells (right). (B) Lung resident memory CD4+ T cell responses. Representative FACS plots (left) and absolute numbers of CD4+ T cells (right) are shown. (C) Heterologous challenges with H3N2 (X-31) virus. Mice previously infected with 50 PFU of H1N1 or H5N2(2:6) virus were challenged with H3N2 (X-31) strain at a dose of 5×10^6^ PFU. The values are expressed as mean ± SD. * denotes statistical significance of <0.05. Data in panels A-B are from two independent experiments pooled together. Data in panel C was performed once.

## Discussion

Infections with avian H5N1 influenza virus induce higher innate immune responses as compared to human H1N1 viruses (21, 36). However, due to inherent differences in replication levels, it is difficult to discern if this hyperactivation of innate immune responses against H5N1 is due to higher viral replication in the lungs. To overcome this caveat, we generated an H5N1 strain sharing the 6 internal genes of H1N1 (H5N1 (2:6)), and observed that the HA and NA of H5N1 can induce higher activation of lung DC. As such, this heightened stimulation of lung DC by H5N1 (2:6) resulted in increased migration of DC to the MLN, and induced robust T cell responses as compared to H1N1 virus. Interestingly, despite the higher numbers of virus specific T cells in the lungs, we observed delayed clearance of H5N1 (2:6) from the lungs of infected mice. This delayed viral clearance correlated with increased levels of PD-1 expression and IL-10 production by CD8+ T cells, which likely suppress cytotoxic T cell functions *in vivo*. Importantly, H5N1 (2:6) infection resulted in decreased numbers of tissue resident memory T cells as compared to H1N1 infection. Taken together, our studies demonstrate that hyperactivation of the innate immune system by H5N1 (2:6) results in suppression T cell functions, delayed viral clearance, and decreased numbers of tissue resident memory T cells.

Unlike seasonal influenza viruses, avian H5N1 influenza viruses can cause severe and often fatal disease in healthy individuals (37, 38). H5N1 infection induces uncontrolled activation of the host immune system, with heightened cytokine levels in the lungs as well as massive infiltration of neutrophils, inflammatory monocytes and inflammatory TNFa/iNOS producing (Tip) DC (39). These infiltrating cells have been implicated in the enhanced virulence of avian H5N1 influenza viruses (26, 27, 39). Moreover, *ex vivo* studies show that H5N1 viruses induce higher human DC activation as compared to H1N1 (25). Similarly, our studies with H5N1 (2:6) demonstrated increased activation of murine lung DC as compared to H1N1 virus, further suggesting that the H5N1 HA/NA are sufficient for higher activation of innate immune cells in the lungs (Figure 1E-F). This increased activation of lung DC in H5N1 (2:6) infected mice was not due to differences in viral replication between strains, as both the reassortant H5N1 (2:6) and H1N1 (PR8) strains showed similar levels of viral replication on days 2, 4 and 6 pi (Figure 3F). These results corroborate prior *ex vivo* studies which indicate that viruses with different HA subtypes can differentially activate primary DC and macrophages (40). However, the consequence of higher DC activation *in vivo* to subsequent adaptive immune responses was previously unknown. Our studies demonstrated that H5N1 (2:6) infection stimulated increased migration and accumulation of DC in the MLN, resulting in robust T cell responses in the lungs (Figure 2C-E). Moreover, we observed increased frequencies of cytokine producing T cells in H5N1 (2:6) infected mice as compared to H1N1 infected mice (Figure 3). Together, these data demonstrate that hyperactivation of lung DC results in increased numbers of virus specific T cells in the lungs of H5N1 (2:6) infected mice.

Prior studies demonstrate that the magnitude and the quality of T cell responses determine the efficiency of viral clearance (41). Previously, we have demonstrated that mice deficient in RIG-I or MAVS mounted poor T cell responses against influenza virus as evidenced by decreased numbers of polyfunctional T cells and delayed viral clearance in the lungs (29). In contrast, despite mounting robust T cell responses, H5N1 (2:6) infected mice showed delayed viral clearance in the lungs as compared to H1N1 infected mice (Figure 3F). This delayed viral clearance in H5N1 (2:6) infected mice was likely due to active suppression of cytotoxic T cell functions *in vivo*, as T cells isolated from H5N1 (2:6) infected mice showed efficient cytotoxic activity against NP peptide pulsed splenocytes (Figure 3G). In corroboration, we observed higher levels of inhibitory signals (PD-1 and IL-10) that likely suppress cytotoxic T cell functions *in vivo* and delay viral clearance (Figure 4A and 3F). In our *in vivo* killing assays, H5N1 (2:6) infected mice showed robust killing of viral peptide loaded splenocytes, suggesting that inhibition of T cells may occur by direct suppression by cell-cell contact rather than by the presence of suppressive cytokine milieu. Prior studies indicate that infection with the lethal mouse adapted PR8 strain (H1N1) resulted in higher PD-1 expression on T cells in comparison to the less virulent X-31 (H3N2) reassortant strain (30). Interestingly, our studies show that infection with H5N1 reassortant (2:6) induced higher PD-1 expression in comparison to PR8 (H1N1) (Figure 4A). As H5N1 viruses have been shown to have broad tissue tropism, it is possible that the increased PD-1 expression observed in H5N1 (2:6) infected mice is likely due to antigen persistence and/or prolonged stimulation of T cells. PD-1 interactions with PD-L1 have been demonstrated to suppress cytotoxic CD8+ T cell functions (42-44). PD-L1 expression is induced during viral infection on a variety of cell types including monocytes, DC, macrophages and epithelial cells(43-46). In our analysis of cell types expressing PD-L1, we observed higher PD-L1 expression on Ly6C^hi^ inflammatory monocytes isolated from H5N1 (2:6) infected mice as compared to H1N1 infected mice (Figure 4C). In addition, the numbers of inflammatory monocytes were higher in H5N1(2:6) infected mice as compared to H1N1 infected mice. It should be noted that PD-L1 expression was observed on others cell types as well, yet there was no significant difference in PD-L1 levels between the two groups (data shown for inflammatory DCs; Figure 4C). In a prior study, anti-PD-L1 treatment of PR8 infected mice showed increased virus specific T cells and decreased viral titers (30); however, anti-PD-L1 treatment did not alter disease outcome, suggesting that there may be additional mechanisms for suppression of T cell functions. In agreement, we observed increased expression of the anti-inflammatory cytokine IL-10 in T cells isolated from H5N1 (2:6) infected mice as compared to H1N1 infected mice (Figure 4C-E). Taken together, our data suggest that higher levels of IL-10 production and PD-1/PD-L1 mediated inhibition likely contribute to suppression of T cell functions and consequently results in delayed clearance of H5N1 (2:6) in the lungs.

Upon viral clearance in the lungs, a portion of virus specific T cells differentiate into tissue resident memory T cells, and these T_RM_ cells are critical for providing heterosubtypic immunity (35, 47). Interestingly, we observed decreased numbers of T_RM_ cells in H5N1 (2:6) infected mice as compared to H1N1 infected mice (Figure 5). It should be noted that despite decreased number of T_RM_ cells, we did not oberserve significant differences protection against challenge from a heterologous H3N2 strain. Our future studies will determine if lowered numbers of T_RM_ cells are due defects in differentiation versus maintenance of T_RM_ cells. Prior studies indicate that Transforming growth factor-β (TGFβ) promote maturation of T_RM_ by inducing the upregulation of CD103 expression (35, 48-50). Co-incidently, influenza viral neuraminidase (NA) can convert latent TGFβ into mature TGFβ; however, the NA of H5N1 is unable to activate TGFβ both *in vitro* and *in* vivo (51, 52). It is possible that the decreased numbers of T_RM_ cells in H5N1 (2:6) infected mice may result from lowered levels of TGFβ activation by viral NA. Apart from TGF-β, interleukin-33 (IL-33) and tumor necrosis factor (TNF) are also known to induce T_RM_ cell like phenotypes (CD69+ CD103+) (50, 53-55). Moreover, homeostatic cytokine IL-15 is required for T_RM_ cell differentiation and survival (55). Thus, it is also possible that H5N1 (2:6) infections may alter the levels of other cytokines that are critical for generation and maintenance of T_RM_ cells. Alternatively, sustained inflammation during H5N1 (2:6) infection may negatively regulate T_RM_ differentiation due to higher levels of IFN-β and IL-12 (56). Further studies are needed to determine if decreased TGFβ levels or sustained higher inflammation in the lungs of H5N1 (2:6) infected mice are responsible for the inefficient differentiation of the T_RM_ population.

In conclusion, our study demonstrates that hyperactivation of innate immune cells by H5N1 (2:6) dampens T cells responses and delays viral clearance in the lungs. This is likely due to higher expression of the inhibitory molecule PD-1 on T cells as well as higher production of the anti-inflammatory cytokine IL-10 by T cells in H5N1 (2:6) infected mice. As such, our studies show that suppression of T cell responses may contribute to the protracted viral replication and prolonged illness associated with avian influenza virus infection in humans.

## Experimental Procedures

### Ethics Statement

All studies were performed in accordance with the principles described by the Animal Welfare Act and the National Institutes of Health guidelines for the care and use of laboratory animals in biomedical research. The protocols for performing murine studies were reviewed and approved by the Institutional Animal Care and Use committee (IACUC) at the University of Chicago.

### Cell Lines

Human embryonic kidney cells (293T, ATCC) were maintained in DMEM (Gibco) supplemented with 10% fetal bovine serum (FBS, Denville Scientific) and penicillin/streptomycin (Pen/Strep, 100 units/mL, Corning). Madin-Darby Canine Kidney (MDCK, ATCC) cells were maintained in Minimum Essential Medium (MEM; Lonza) supplemented with 10% FBS and Pen/Strep (100 units/mL).

### Viruses

The generation of H1N1-GFP (A/Puerto Rico/8/1934) has been described earlier(57). H5N1-GFP (A/Vietnam/1203/2004; low pathogenic without the multibasic site in HA), which contains a GFP reporter in the NS segment, was generated following a similar protocol(57). H5N1 (2:6) (A/Vietnam/1203/2004; low pathogenic without the multibasic site in HA), which contains the 6 internal genes from the PR8 strain, was rescued using standard reverse genetics techniques (57, 58). Briefly, 0.5 μg of each of the six pDZ plasmids representing PB2, PB1, PA, NP, NS and M from A/Puerto Rico/8/1934 (PR8) and two pPol-I plasmids representing the HA (low pathogenic) and NA segments of H5N1 were transfected into a cell mixture containing 293T-MDCK using Lipofectamine 2000 (Invitrogen). After 48hrs, 200µl of the rescue supernatants was used to infect fresh MDCK cells seeded in 6 well plates. The successful rescue of recombinant viruses was confirmed by performing hemagglutination assay with chicken red blood cells. After plaque purification, the recombinant viruses were amplified in 10-day old specific pathogen free eggs (Charles River). Viral titers were determined by plaque assay in MDCK cells using standard techniques.

### Mice infection

C57BL/6 mice were purchased from Jackson Laboratory and bred in specific pathogen free (SPF) facilities maintained by the University of Chicago Animal Resource Centre. All experiments were performed with gender-matched mice of 6-8 weeks of age. For influenza virus infections, mice were anesthetized with ketamine/xylazine (i.p 80/10mg/kg) and infected intranasally with the indicated dose of virus diluted in 25 μl of PBS.

### Quantitative RT-PCR analysis

Total RNA from lung tissue was extracted using Trizol (Life technologies) following the manufacturer’s instructions, and cDNA was synthesized with SuperScript II using Oligo dT primers (Roche Diagnostics). Quantitative PCR was performed using previously described gene specific primers in an ABI7300 Real Time PCR system with SYBR Green Master Mix (Invitrogen) (12).

### Flow cytometric analyses

Preparation of lung samples for flow cytometric analysis and T cell assays were performed following techniques previously described by us (29). Briefly, after euthanization, murine lungs were perfused with 10 ml of PBS, excised and finely chopped with scissors, and digested in 0.4mg of Collagenase in HBSS/10%FBS for 45 minutes at 37C. Mediastinal lymph nodes (MLN) were carefully isolated and digested in 0.2mg of collagenase in HBSS/10%FBS for 15 minutes at 37C. To prepare single cell suspensions, collagenase treated lung tissues and MLN were passed through a 19G blunt needle a few times and filtered through a 70µm cell strainer. After two washes in FACS buffer (PBS containing 1% FBS and 2mM EDTA), the cells were subjected to RBC lysis (Biowhitaker) for 3 minutes followed by two washes with FACS buffer. The single cell preparations were resuspended in FACS buffer containing 10µg/ml Fc receptor block and incubated for 15 minutes. For DC subset analysis, lymph node and lung cells were stained with antibodies against multiple surface antigens: anti-CD45 (2µg/ml, 30-F11; Biolegend), anti-SiglecF (1µg/ml, E50-2440; BD Biosciences), anti-CD11c (2µg/ml, N418; Biolegend), anti-MHC-II (2µg/ml, M5/114.15.2; Biolegend), anti-CD103 (2µg/ml, 2E7; eBiosciences), anti-CD11b (1µg/ml, M1/70; Biolegend), anti-CD86 (2µg/ml, GL-1; Biolegend), anti-Ly6G (1µg/ml, 1A8; Biolegend), anti-Ly6C (2µg/ml HK1.4; Biolegend), anti-CD4 (2µg/ml, RM4-4; Biolegend), anti-CD3 (2µg/ml, 145-2C11; eBiosciences), and anti-CD8 (1µg/ml, 53-6.7; eBiosciences). Dead cells were stained with Live/Dead Fixable Near IR Staining Kit (Life Technologies) in PBS for 15 minutes on ice. Surface stained samples were fixed with FACS buffer containing 0.1% formaldehyde and analyzed using the BD LSR-II flow cytometer. Data analysis was performed using FlowJo software (Treestar Corp.).

### DC and T cell assays

Bone marrow derived dendritic cells (BMDC) were generated from C57BL/6 mice and T cell re-stimulation experiments were performed as previously described (59), (60). Briefly, BMDC were infected with X-31 (H3N2) at an MOI of 0.5 for 5h, washed with PBS 3 times to remove unbound virus, and resuspended in Iscove’s Modified Dulbecco’s Media (IMDM) with 10% FBS (Invitrogen). T cells from the lungs of naïve or infected mice (day 8 pi) were enriched using the Pan T cell Isolation Kit II (Miltenyi Biotec) and co-cultured with infected BMDC at a ratio of 10:1 for 2-3 hours followed by the addition of Brefeldin A (5µg/ml; eBiosciences). The cells were further incubated for an additional 8-10h at 37C. Ex vivo peptide stimulation studies were performed using MHC-I NP_366-374_ (ASNENMETM) or MHC-II restricted NP_311–325_ (QVYSLIRPNENPAHK) peptides. The cells were first stained for cell surface markers as described above, followed by intracellular staining for cytokines. For intracellular staining, cells were incubated in Permeabilization and Fixation buffer (BD Pharmingen) for 45 minutes followed by 2 washes in a PBS buffer containing 1% FBS and 0.5% Saponin (Sigma, St Louis, MO). Intracellular staining for anti-IFNγ (2µg/ml, XMG1.2; Biolegend), anti-Granzyme B (2µg/ml, GB11; Biolegend), and IL-10 (2µg/ml, JES5-16E3; Biolegend) was performed on ice for 30 minutes.

For T cell tetramer staining, lymphocytes from the lungs of influenza virus infected mice were enriched using Ficoll-Hypaque (GE Healthcare Life Sciences) density gradient and stained with H-2D^b^ restricted tetramers conjugated to fluorophore R-phycoerythrin (PE) (NP_366-374_ ASNENMETM or PA_224-233_ SSLENFRAYV).

### In vivo killing assay

Single cell suspension was prepared from mice spleen and the cells were pulsed either with 1μM NP_366-374_ peptide or OT-1 peptide (Ova _257-264_) for 1hour and labelled with 1μM CFSE (CFSE^low^) or 5 μM CFSE(CFSE^high^) respectively following manufacturer’s instructions (Life Technologies). A mixture of 2×10^6^ CFSE^low^ and CFSE^high^ splenocytes were intravenously injected to gender matched naïve mice or the mice which had been intranasally infected 8 days ago with H1N1 or H5N1(2:6) virus. After 5 hours of injection, mice splenocytes were analysed for CFSE positive cells by flow cytometry. Percent killing was determined using the following equation:

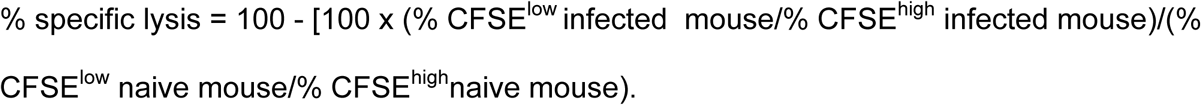

### Analysis of tissue resident memory T cells

Lung resident memory CD8+T cells were analyzed as previously described(61). Mice were intranasally infected with H1N1 or H5N1 (2:6) virus. On day 30 post infection, mice were intravenously injected with 1μg anti CD8β antibody 5 minutes before tissue harvest. Lung tissues were perfused with PBS and single cell suspension were prepared after digestion with collagenase as described before. Cells were blocked first with FcRγIII/II antibody and stained with H-2D^b^ restricted tetramer conjugated to fluorophore R-phycoerythrin (PE) (NP_366-374_ ASNENMETM). Tetramer labelled cells were washed and stained with anti-CD4 (2µg/ml, RM4-4; Biolegend), anti-CD3 (2µg/ml, 145-2C11; eBiosciences), anti-CD8a (1µg/ml, 53-6.7; eBiosciences), anti-CD44 (IM7; Biolegend), anti-CD103 (2µg/ml, 2E7; eBiosciences) and anti-CD69 (H1.2F3; Biolegend). Dead cells were stained with Live/Dead Fixable Near IR Staining Kit (Life Technologies) in PBS for 15 minutes on ice. Surface stained samples were fixed with FACS buffer containing 0.1% formaldehyde and analyzed using the BD LSR-II flow cytometer. Data analysis was performed using FlowJo software (Treestar Corp.)

### Statistical analysis

Data was analyzed using Prism GraphPad software and statistical significance was determined by one-way ANOVA or the unpaired Student’s t test. *, **, *** denotes significance of <0.05, <0.01, <0.001, respectively; ns denotes not significant.

## Author Contributions

MK, BM and SM conceived and designed the study. MK, KF and SM performed experiments. MK, JTP, SM and BM wrote the manuscript. All authors approved the manuscript.

## Acknowledgements

We are grateful to Dr. Adolfo Garcia-Sastre at the Icahn School of Medicine for providing numerous reagents used in this study. H5N1 reverse genetics plasmids were kindly provided by Dr. John Steel at Emory University. We would like to thank the NIH Tetramer Core Facility at Emory University for providing us with influenza virus specific T cell tetramers. We would also like to thank the staff at the Office of Research Safety and Animal Resource Center at the University of Chicago for their excellent support.

